# Shouldering the challenge of deciphering avian palate evolution

**DOI:** 10.1101/2025.06.10.658509

**Authors:** Juan Benito, Pei-Chen Kuo, Christopher R. Torres, Guillermo Navalón, Olivia Plateau, Alexander D. Clark, Elizabeth M. Steell, Daniel J. Field

## Abstract

Wilken et al. (1) investigate the evolution of avian palatal kinesis using comparative morphology and biomechanical modelling. While the study’s topic and approach are timely, its conclusions are marred by inadequate taxon sampling and morphological misinterpretations.

## Main text

Reinterpreting the *Janavis* pterygoid (2) as a juvenile bird coracoid (1) is contradicted by clear morphological and developmental evidence (Fig. 1). The concave, ossified ‘sternal articulation’ of the would-be coracoid (1) is unossified in juvenile bird coracoids, yet resembles the quadrate articulation of the pterygoid in several extant taxa (Fig. 1B). Similarly, *contra* (1), the basipterygoid articulation in *Janavis* closely resembles that of several extant neognathous birds in its form and position (Fig. 1C).

**Fig. 1.**
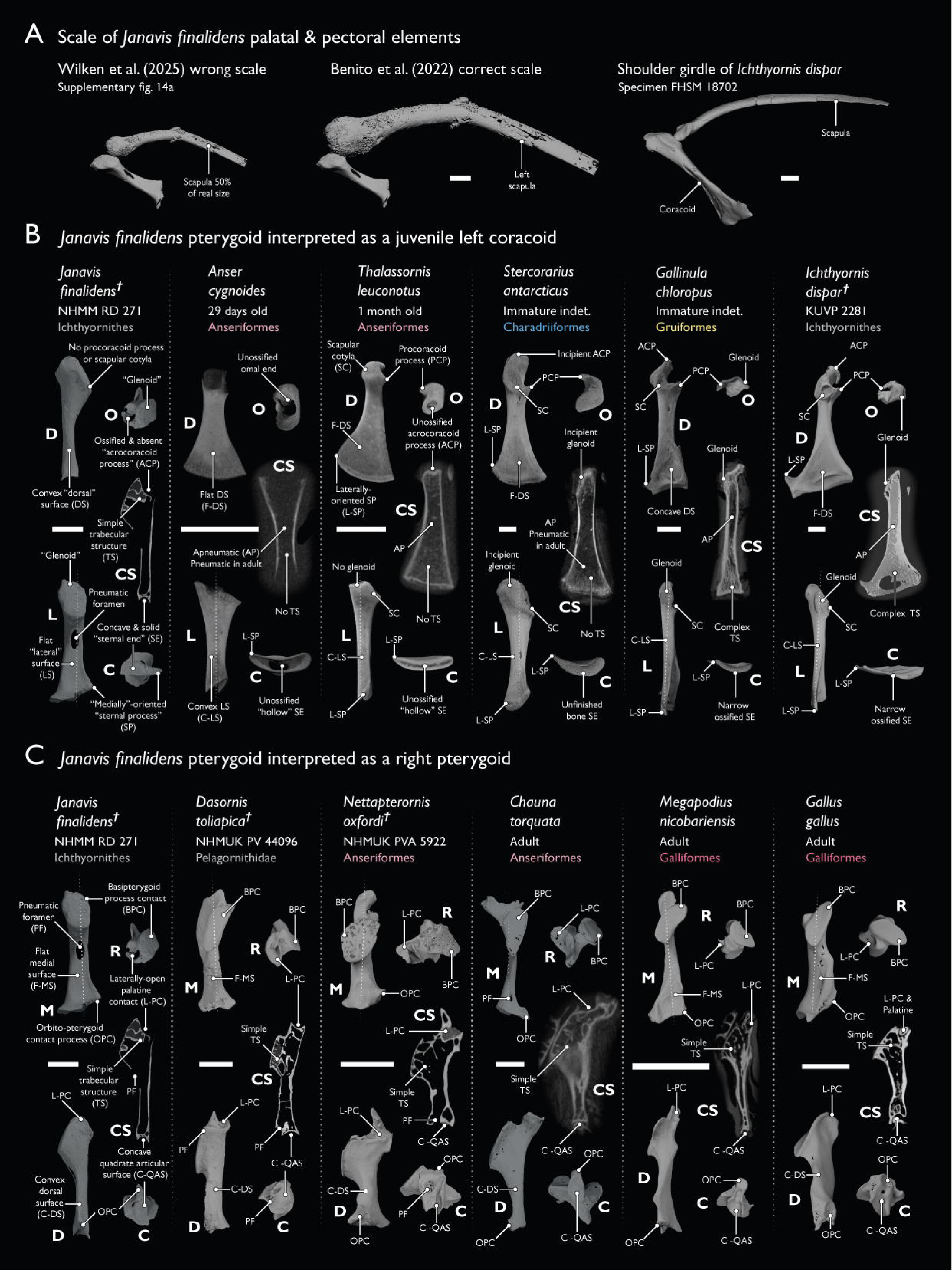
The pterygoid of *Janavis finalidens* is not a juvenile coracoid. **A)** The scapula of *Janavis* was downscaled to 50% of its real size by (1) (see their Fig. S14). When correctly scaled (Fig. 1a and Extended Data Fig 2b-d in (2)) the pterygoid and scapula do not match (nor does the pterygoid resemble a coracoid in the first place—see main text). **B)** If interpreted as an immature coracoid, the pterygoid of *Janavis* differs from all known avian coracoids in: 1) glenoid development ontogenetically preceding the scapular cotyle, procoracoid, and acrocoracoid; 2) exhibiting a ‘dorsally’ convex and ‘laterally’ flat shaft; 3) the authors’ (1) ‘sternal process’ (presumably the sternolateral process) being medially, rather than laterally, oriented; 4) exhibiting fully ossified concave surfaces on both omal and sternal ends, which are instead open and hollow or formed by unfinished bone in immature bird coracoids (the sternal end of the coracoid is only concave in adult, highly derived Apodiformes); 5) being highly pneumatic with a simple trabecular structure, as pneumaticity only develops late in post-hatching development in extant birds. **C)** If correctly interpreted as a right pterygoid (*contra* (2), where it was considered a left pterygoid), all features of the element are found in fossil and extant neognathous bird pterygoids, including: 1) a large, medially-oriented basipterygoid process facet reaching the rostral end (present in Galliformes, *contra* (1)); 2) a laterally and often dorsally open, cup-shaped palatine contact (present across Galloanserae); 3) a tubular shaft that is medially flat and dorsally convex (present in Galliformes and Pelagornithidae); 4) a simple, concave quadrate articular surface (present in Anseriformes); 4) a marked orbito-pterygoid contact process (present in Galliformes and some Anseriformes); 5) a heavily pneumatised internal structure with simple trabeculae (present across Galloanserae). Bold letters indicate anatomical views: caudal (C), dorsal (D), lateral (L), medial (M), rostral (R). Cross-sectional (CS) planes are indicated with dotted lines over each element. All scale bars equal 5 mm.

The authors identify a supposed glenoid and acrocoracoid on the purported coracoid (1), yet in developing bird coracoids these structures only arise after the scapular cotyle, which is absent in the fossil. Importantly, the positions of these structures do not shift relative to one-another throughout extant bird ontogeny, which would be necessary for the element to come to resemble a coracoid (Fig. 1B). Since ichthyornithine postcranial morphology and growth generally fall within the range of variation of extant birds (3), there is no reason to think these would be any different in *Janavis*. Crucially, the authors of (1) must have reduced the size of the *Janavis* scapula by ~50% to ‘match’ it with the putative coracoid (see their Supplementary Fig. 14A; our Fig. 1A), yet this is unmentioned.

The above issues notwithstanding, the ichthyornithine hemipterygoid alone—with an articular condyle like those of prokinetic neognaths —suggests a mobile intrapterygoid joint (Fig. 2A). Wilken et al. seemingly agree (“…*Ichthyornis*… possesses a segmented, potentially propulsive pterygoid…” (1)); yet, they interpret palatal kinesis as a neognath autapomorphy by asserting that stem bird musculature was incompatible with palate mobility. This rests upon misinterpretations of the functional correlates of quadrate morphology. First, elongate orbital processes in *Ichthyornis* and *Hesperornis* are declared to impose constraints on kinesis due to articulation with the neurocranium (1). However, many neognaths exhibit equivalently long (or longer) orbital processes closely approaching the neurocranium without precluding prokinesis (Fig. 2B; (5); https://doi.org/10.5281/zenodo.15619495). Second, *contra* (1), the bicondylar nature of the *Ichthyornis* quadrate would not have precluded coupled palatal kinesis, as evidenced by bicondylar Anseriformes (Fig. 2B) (6). Further morpho-functional misconceptions underpin problematic cranial linkage models in (1) (Fig. 2C), yet the authors’ own ancestral reconstructions of cranial linkages and protractor force seemingly support palatal mobility as plesiomorphic for all crown birds (their Figs. 3A, 4), rather than being apomorphic for Neognathae as they argue.

**Fig. 2.**
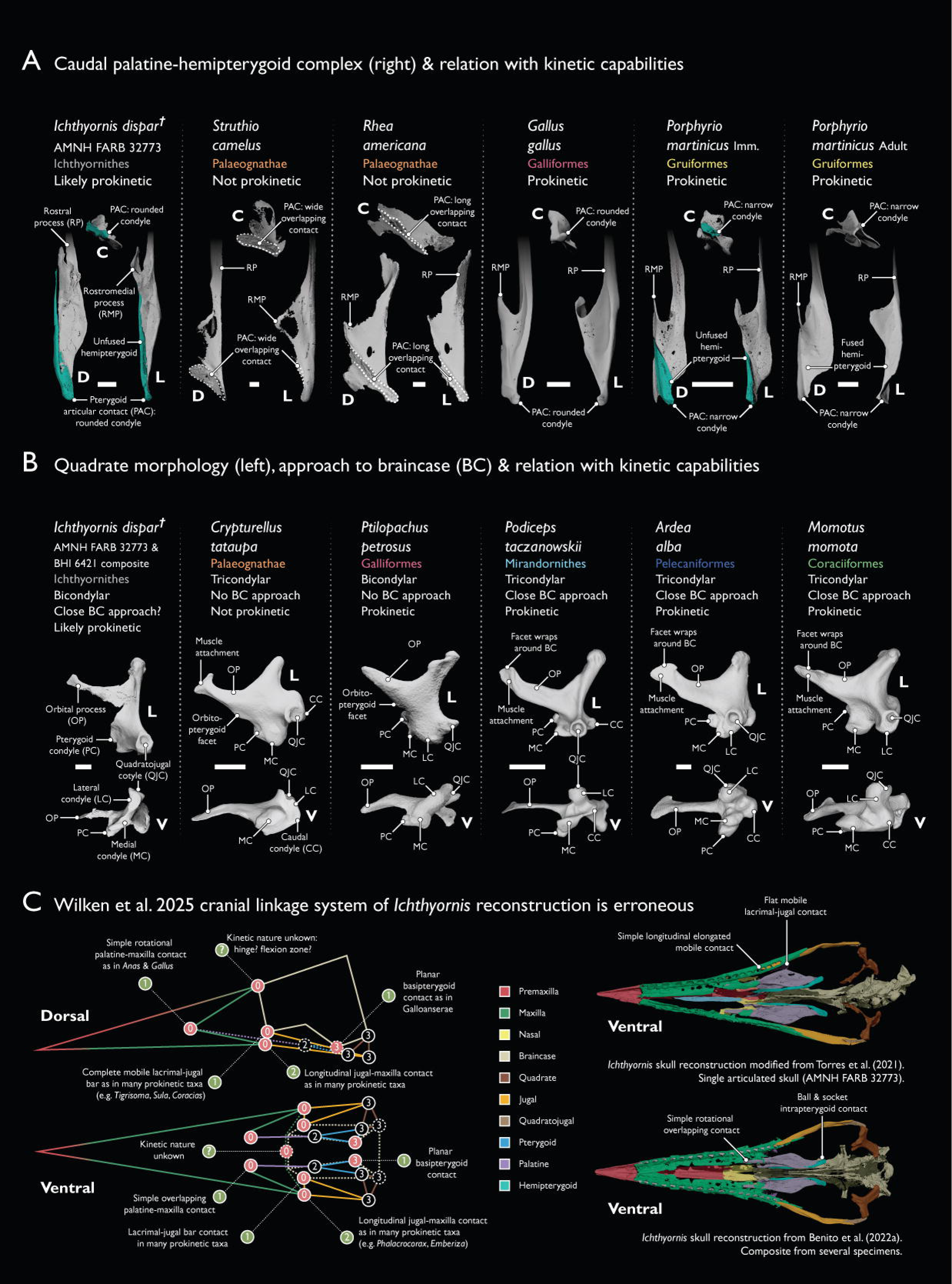
Morphology of the suspensorial system in Ichthyornithes suggests prokinetic capabilities. **A)** *Ichthyornis* exhibited a mobile intrapterygoid (hemipterygoid-pterygoid) articulation, like the condition in unsegmented immature (imm.) neognaths (hemipterygoid highlighted in teal shade). The condylar morphology of the pterygoid articular contact resembles that of prokinetic Galliformes and Neoaves, and contrasts with the long or wide overlapping contacts of palaeognaths. Long rostral processes of the palatine are obscured for clarity. **B)** The quadrates of *Ichthyornis* and *Hesperornis* (bicondylar, with long orbital processes interpreted as contacting the braincase (1)) would not preclude palatal kinesis. Both features are present across a wide range of prokinetic neognath taxa unsampled by (1). This includes taxa where the orbital process 1) wraps around and closely approaches the neurocranium; 2) terminates in an expansion whose lateral surface anchors the M. pseudotemporalis profundus; and 3) is presumably associated with depressions or ‘cotylae’ on the braincase (e.g., *Ardea*; see https://doi.org/10.5281/zenodo.15619495). **C)** The ichthyornithine linkage model proposed by (1) enforces immobility at contacts between bones (e.g., the jugal and the lacrimal) which do not preclude prokinesis in extant birds. We present corrected linkage values following the scheme of (1), indicated in green, and illustrate the support for these from previous reconstructions of the skull of *Ichthyornis*. Bold letters indicate anatomical views: caudal (C), dorsal (D), lateral (L), ventral (V). All scale bars equal 2.5 mm. Digitally segmented *Struthio camelus* palatine in A courtesy of Annabel K. Hunt.

Contemporary research in avian macroevolution seeks to evaluate associations between brain and skull transformations (7-9); however, this link is weakly substantiated by (1), as several taxa with palatal kinesis (e.g., Anseriformes) share plesiomorphically low degrees of encephalization with akinetic palaeognaths and stem birds (10).

## Conclusions

The origin of palatal kinesis was a key milestone in avian evolutionary history—careful morphological interpretation will be crucial to clarify how this major transformation occurred and influenced avian evolutionary history.

