## Supplementary Information for "Shouldering the challenge of deciphering avian palate evolution"

Data hosted in the Zenodo Project “Benito et al., Shouldering the challenge of deciphering avian palate evolution” accessible at https://doi.org/10.5281/zenodo.15619496).

This project contains the data supporting Benito et al., response to Wilken et al. (2025), including morphological observations, 3D mesh models of relevant skeletal elements and videos of dissections exploring avian cranial kinesis.

- “Specimen list – Morphosource links”: this file contains a list of all the specimens figured in Figs 1 and 2, including details on taxonomy, museum catalog numbers and relevant elements studied. Meshes for all elements and relevant CT datasets are housed in the online repository Morphosource, links for each individual element are provided here.
- “Quadrate_orbital_process_kinesis_dataset”: this spreadsheet contains a dataset of morphological observations exploring the relation between specific morphologies (e.g. an elongated quadrate orbital process) and the presence of cranial kinesis. It also characterises the presence of cranial kinesis (including both prokinesis and rhynchokinesis) in all surveyed taxa, including published sources and personal observations from osteological and dissection specimens.
- “Quadrate mobility dissection videos”: this folder contains a series of videos of dissections exploring cranial kinesis in freshly deceased birds. A key “Dissection videos key” is provided detailing each video.
- “Specimen STLs”: this folder (and subfolders” contain mesh models (in .stl format) for all skeletal elements figured in Figs 1 and 2. These are divided per element (coracoid, pterygoid, palatine and quadrate) and those of Janavis are provided in their own subfolder. Links to all these elements housed in Morphosource are provided in the specimen list above.
