## Supplementary dataset 1 for "Shouldering the challenge of deciphering avian palate evolution"

All data available at Morphosource Project ID 000748651 - Accessible at https://www.morphosource.org/projects/000748651?locale=en

| Clade | Genus | species | Specimen | Type of specimen | Element | Data type | Morphosource link |
|---|---|---|---|---|---|---|---|
| Ichthyornithes | Ichthyornis | dispar | FHSM 18702 | Fossil | Coracoid | Mesh | ark:/87602/m4/405003 |
| Ichthyornithes | Ichthyornis | dispar | FHSM 18702 | Fossil | Scapula | Mesh | ark:/87602/m4/404925 |
| Ichthyornithes | Ichthyornis | dispar | KUVP 2281 | Fossil | Coracoid | Mesh | ark:/87602/m4/405518 |
| Ichthyornithes | Ichthyornis | dispar | BHI 6421 | Fossil | Quadrate | Mesh | ark:/87602/m4/509698 |
| Ichthyornithes | Ichthyornis | dispar | AMNH FARB 32773 | Fossil | Quadrate | Mesh | ark:/87602/m4/748948 |
| Ichthyornithes | Ichthyornis | dispar | AMNH FARB 32773 | Fossil | Palatine right | Mesh | ark:/87602/m4/748957 |
| Ichthyornithes | Ichthyornis | dispar | AMNH FARB 32773 | Fossil | Palatine left | Mesh | ark:/87602/m4/748954 |
| Ichthyornithes | Ichthyornis | dispar | AMNH FARB 32773 | Fossil | Hemipterygoid | Mesh | ark:/87602/m4/748951 |
| Ichthyornithes | Ichthyornis | dispar | AMNH FARB 32773 | Fossil | Braincase | Mesh | ark:/87602/m4/748963 |
| Ichthyornithes | Ichthyornis | dispar | AMNH FARB 32773 | Fossil | Various elements for reconstruction | Mesh (various) | ark:/87602/m4/367065 |
| Ichthyornithes | Janavis | finalidens | NHMM RD 271 | Fossil | Pterygoid | CT Data | ark:/87602/m4/445881 |
| Ichthyornithes | Janavis | finalidens | NHMM RD 271 | Fossil | Pterygoid | Mesh | ark:/87602/m4/469732 |
| Ichthyornithes | Janavis | finalidens | NHMM RD 271 | Fossil | Pterygoid rostral | Mesh | ark:/87602/m4/469729 |
| Ichthyornithes | Janavis | finalidens | NHMM RD 271 | Fossil | Pterygoid caudal | Mesh | ark:/87602/m4/469726 |
| Ichthyornithes | Janavis | finalidens | NHMM RD 271 | Fossil | Fossil block (includes scapula) | CT Data | ark:/87602/m4/445292 |
| Ichthyornithes | Janavis | finalidens | NHMM RD 271 | Fossil | Scapula | Mesh | ark:/87602/m4/470249 |
| Palaeognathae | Struthio | camelus | UMZC Uncatalogued | Osteological | Palatine | Mesh | ark:/87602/m4/749087 |
| Palaeognathae | Rhea | americana | FMNH:Birds:339616 | Osteological | Palatine | Mesh | ark:/87602/m4/748960 |
| Palaeognathae | Crypturellus | tataupa | UMMZ:birds:201948 | Osteological | Quadrate | Mesh | ark:/87602/m4/550328 |
| Pelagornithidae | Dasornis | toliapica | NHMUK PV 44096 | Fossil | Pterygoid | CT Data | ark:/87602/m4/748933 |
| Pelagornithidae | Dasornis | toliapica | NHMUK PV 44096 | Fossil | Pterygoid | Mesh | ark:/87602/m4/470232 |
| Anseriformes | Anser | cygnoides | UMZC Uncatalogued | Immature (29 days old) - frozen | Pectoral region | CT Data | ark:/87602/m4/748805 |
| Anseriformes | Anser | cygnoides | UMZC Uncatalogued | Immature (29 days old) - frozen | Coracoid | Mesh | ark:/87602/m4/748917 |
| Anseriformes | Thalassornis | leuconotus | UMZC Uncatalogued | Immature (1 month old) - frozen | Pectoral region | CT Data | ark:/87602/m4/748817 |
| Anseriformes | Thalassornis | leuconotus | UMZC Uncatalogued | Immature (1 month old) - frozen | Coracoid | Mesh | ark:/87602/m4/748911 |
| Anseriformes | Nettapterornis | oxfordi | NHMUK PVA 5922 | Fossil | Pterygoid | CT Data | ark:/87602/m4/748941 |
| Anseriformes | Nettapterornis | oxfordi | NHMUK PVA 5922 | Fossil | Pterygoid | Mesh | ark:/87602/m4/470238 |
| Anseriformes | Chauna | torquata | UMZC Uncatalogued | Frozen | Head | CT Data | ark:/87602/m4/749314 |
| Anseriformes | Chauna | torquata | NHMUK s/2012.31.1 | Osteological | Pterygoid | Mesh | ark:/87602/m4/470176 |
| Galliformes | Megapodius | nicobariensis | UMZC 14/Meg/f/3 | Skin | Head | CT Data | ark:/87602/m4/749104 |
| Galliformes | Megapodius | nicobariensis | UMZC 14/Meg/f/3 | Skin | Pterygoid | Mesh | ark:/87602/m4/461880 |
| Galliformes | Gallus | gallus | UMZC:Vertebrates:400 | Osteological | Skull | CT Data | ark:/87602/m4/749581 |
| Galliformes | Gallus | gallus | UMZC:Vertebrates:400 | Osteological | Pterygoid | Mesh | ark:/87602/m4/749581 |
| Galliformes | Gallus | gallus | UMZC:Vertebrates:400 | Osteological | Palatine | Mesh | ark:/87602/m4/749728 |
| Galliformes | Ptilopachus | petrosus | NHMUK 1968.9.2 | Spirit specimen | Quadrate | Mesh | ark:/87602/m4/509841 |
| Mirandornithes | Podiceps | taczanowskii | UMMZ:birds:156782 | Osteological | Quadrate | Mesh | ark:/87602/m4/551340 |
| Gruiformes | Gallinula | chloropus | UMZC Uncatalogued | Immature (age indet.) - frozen | Pectoral region | CT Data | ark:/87602/m4/748838 |
| Gruiformes | Gallinula | chloropus | UMZC Uncatalogued | Immature (age indet.) - frozen | Coracoid | Mesh | ark:/87602/m4/748905 |
| Gruiformes | Porphyrio | martinicus | UMZC 15/Ral/43/b1/1 | Skin | Palatine | Mesh | ark:/87602/m4/748968 |
| Gruiformes | Porphyrio | martinicus | UMZC Uncatalogued | Immature (Age indet.) - spirit | Palatine | Mesh | ark:/87602/m4/749755 |
| Gruiformes | Porphyrio | martinicus | UMZC Uncatalogued | Immature (Age indet.) - spirit | Hemipterygoid | Mesh | ark:/87602/m4/749748 |
| Charadriiformes | Stercorarius | antarcticus | UMZC Uncatalogued | Immature (age indet.) - frozen | Pectoral region | CT Data | ark:/87602/m4/748830 |
| Charadriiformes | Stercorarius | antarcticus | UMZC Uncatalogued | Immature (age indet.) - frozen | Coracoid | Mesh | ark:/87602/m4/748908 |
| Pelecaniformes | Ardea | alba | UMZC:Vertebrates:338.H | Osteological | Quadrate | Mesh | ark:/87602/m4/749665 |
| Coraciiformes | Momotus | momota | NHMUK Zoo S/2015.23.4 | Osteological | Quadrate | Mesh | ark:/87602/m4/551250 |
